# Chronic Sleep Disruption Activates the Transcriptional Unfolded Protein Response and Induces Compartment-Specific Proteomic Changes in Mice

**DOI:** 10.64898/2026.11.03.763923

**Authors:** Claudia Espinosa-Garcia, Dilpreet Kour, Upasna Srivastava, Sneha Malepati, Prateek Kumar, Samira Parhizkar, Brendan Tobin, Wooyoung Eric Jang, Rashmi Kumari, Britney Campbell, Jacob Stagray, Maria Henley, Xinran Liu, Levi B Wood, Victor Faundez, Srikant Rangaraju

## Abstract

Sleep disruption (SD) is associated with adverse brain health and increased risk of dementia, including Alzheimer’s disease (AD), yet the pathological mechanisms underlying this association are not fully understood. Preclinical studies have shown that acute SD impairs cognition by altering vulnerable synapses (molecular composition, ultrastructure, and function). However, the cumulative effects of chronic SD on the adult brain and its vulnerable synapses, that might contribute to AD risk, remain poorly defined. Here, we investigated the impact of chronic SD on brain transcriptomic signatures and subcellular compartment-specific proteomic profiles to identify genes, proteins, and signaling pathways that might confer AD risk. Chronic SD induced a modest transcriptional stress response globally, whereas alterations were more pronounced in the synaptosomal (synapse-enriched P2 fractions) and whole-brain proteomes. Remarkably, chronic SD elicited unique proteomic alterations at the synaptic compartment were distinct from whole-brain effects, including increased proteins involved in synaptic vesicle endocytosis, regulation of dendritic spine morphogenesis, and regulation of long-term synaptic depression, compared to increased proteins involved in immune/inflammatory, glial responses and neuronal function at the whole-brain level. Consistent with the chronic SD-induced molecular signatures, we observed subtle ultrastructural changes in vulnerable CA1 hippocampal synapses, including reduced pre-synapse size and decreased synaptic vesicle density, which suggest that chronic SD modifies synaptic transmission and plasticity. Together, our findings indicate that chronic SD differentially impacts the brain and the synaptic proteome, nominating candidate therapeutic targets and signaling pathways for future mechanistic studies of chronic SD and how SD contributes to increased AD risk.

**Highlights:**

- Brain transcriptomics reveals chronic SD activates the unfolded protein response.
- Compartment-specific proteomics shows chronic SD drives unique compensatory changes.
- Chronic SD subtly alters the ultrastructure of vulnerable hippocampal CA1 synapses.
- We nominate candidate pathways that link effects of chronic SD to AD risk.

## INTRODUCTION

Sleep is essential for brain health, supporting key cognitive processes (e.g., synaptic plasticity and memory consolidation) (1–3), and enabling the clearance of neurotoxic metabolic products (e.g., amyloid-β and tau) via pathways that include the glymphatic system (4, 5). Thus, sufficient and high-quality sleep should be considered a health priority (6–9). However, modern lifestyles contribute to sleep disruption (SD). According to the World Health Organization, SD affects millions of people globally, and is increasingly recognized as a major public health concern associated with the development of chronic diseases (8, 10–12). In the United States, the most recent survey from the National Center for Health Statistics indicates that 30.5% of adults experience short sleep duration (< 7 h of sleep per day) and persistent sleep difficulties (13).

Strong evidence from human studies supports chronic SD as a risk factor for the development and progression of dementia, including Alzheimer’s disease (AD) (8, 14–23). For instance, disruptions in sleep architecture (e.g., reduced sleep efficiency, altered sleep duration, and diminished time spent in specific sleep stages) are associated with increased overnight Aβ production (24), greater Aβ accumulation in the brain (25), and mild cognitive impairment (26). Despite the strong association between chronic SD and AD pathology, the exact molecular mechanisms by which chronic SD impacts the brain remain unclear. Synapses are particularly vulnerable to sleep loss (27, 28), highlighting the need to examine synapse-specific changes in response to chronic SD. Preclinical studies show that acute SD impairs cognition by altering vulnerable synapses at the level of their molecular composition, ultrastructure, and function (27, 29–33). However, critical gaps remain in our understanding of how chronic SD drives cumulative synaptic alterations in the brain, with important implications for developing therapeutic interventions that protect vulnerable synapses and prevent cognitive impairment.

To identify genes, proteins, and signaling pathways altered by the cumulative effects of chronic SD in the brain and at the level of synapses, we performed bulk transcriptomic profiling of adult mouse brain and proteomic analysis of crude synaptosomes (synapse-enriched P2 fractions). Bulk brain transcriptomics revealed modest gene changes induced by chronic SD, mainly involving cellular stress and the unfolded protein response. In contrast to modest bulk brain transcriptome changes, chronic SD induced pronounced alterations in the synaptosomal (P2 fraction) and the whole-brain (homogenate) proteomes. Remarkably, gene set variation analysis (GSVA) identified differential effects of chronic SD on molecular pathways in the synaptic compartment compared to the whole-brain, indicative of unique vulnerability mechanisms of the synapse to chronic SD. These molecular signatures were accompanied by subtle ultrastructural changes in vulnerable CA1 hippocampal synapses, including reduced pre-synapse size and decreased synaptic vesicle density, which suggest that chronic SD modifies synaptic transmission and plasticity. Collectively, these results nominate molecular candidates and signaling pathways implicated in chronic SD-associated synaptic dysfunction and increased AD risk, highlighting potential targets for therapeutic intervention.

## EXPERIMENTAL PROCEDURES

### Animals

All animal procedures were performed in accordance with the National Institutes of Health Guide for the Care and Use of Laboratory Animals with the approval of the Yale University Institutional Animal Care and Use Committee (IACUC Protocol #2023-20536). The experiments reported here are in accordance with the ARRIVE guidelines. Male wild-type (WT) C57BL/6J mice were obtained from The Jackson Laboratory (Strain #000664) at an age of 8 weeks and group-housed under standard temperature and humidity conditions (22 ± 2 °C) on a 12:12 h light/dark cycle (lights on at 07:00 [Zeitgeber time 0, ZT0] and off at 19:00 [ZT12] (34)). Food and water were provided *ad libitum*.

### Experimental Design and Statistical Rationale

Given that the molecular consequences of chronic SD, particularly those events occurring at vulnerable synapses, remain poorly defined, this study aims to identify genes, proteins, and signaling pathways altered by chronic SD in the brain and its synaptic compartment. At 6.5 months of age, mice randomly assigned to either the SD group or the undisrupted sleep control group. A sample size of N=10 mice per group was selected based on our previous experience with omics studies (35–38) and established protocols using chronic SD paradigms (39–42). Mice were subjected to a chronic SD paradigm for 6 h/day, from ZT3 to ZT9, over 6 consecutive weeks, or to undisrupted sleep. The next day (18 h recovery sleep) after completing the 6-week chronic SD paradigm, mice were euthanized between ZT3 and ZT5 to minimize circadian influence on molecular signaling, gene expression, protein synthesis and synaptic structure (32, 39), with alternated sampling between SD and control mice. A subset of mice (N=7/group) was deeply anesthetized with isoflurane, and decapitated, their brains were rapidly collected and used for omics analysis. On the one hand, bulk brain tissue samples were immediately flash-frozen in liquid nitrogen and stored at -80°C until processing, total RNA was isolated and further analyzed by RNAseq to identify differentially-expressed genes (DEGs). On the other hand, total brain homogenates and crude synaptosomes (P2 fractions) were freshly prepared from the same animals by differential centrifugation and analyzed by label-free quantitative mass spectrometry (LFQ‒MS) to identify differentially-enriched proteins (DEPs) and over-represented biological pathways. Western blot (WB) and transmission electron microscopy (TEM) were used for quality assessment of P2 fractions. Additionally, homogenates and P2 fractions were assayed using multiplexed Luminex to measure chronic SD-induced alterations in phospho-proteins of the MAPK and the mTOR pathways. Another subset of mice (N=3/group) was deeply anesthetized with isoflurane and transcardially perfused with PBS followed by 4% paraformaldehyde. Fixed brains were dissected in 2 hemispheres and used for histological assessments, i.e., Nissl staining, immunofluorescence, and TEM. Plasma samples were collected at euthanasia to quantify the levels of corticosterone (main stress hormone in rodents (43)) and neurofilament light chain (NfL, a biomarker of axonal injury in neurodegenerative diseases (44)) using ELISA.

### Chronic SD Paradigm

Prior to starting the chronic SD paradigm, mice were transferred to a separated room, randomly assigned to either the SD group or the undisrupted sleep control group, and allowed to acclimate to the animal facility and housing conditions for one week. For the chronic SD paradigm, mice were group-housed in automated sleep fragmentation chambers (LaFayette Instruments; 80391) equipped with a bar set to move every 30 sec for 6 h/day, from ZT3 to ZT9, over 6 consecutive weeks as previously reported (39). These chambers have been validated and widely used in previous studies to model chronic SD in mice (39, 45–49), with the advantage that group housing reduces social isolation stress. SD mice were monitored daily for signs of distress or health decline. For the control group, a second set of mice were moved into similar home cages without the swipe bar, and group-housed alongside SD group to account for environmental stressors (e.g., noise and handling). Body weight was recorded weekly for 8 weeks, from the start of acclimation period through euthanasia. Percentage body weight gain was calculated as (Wt/Wi) x 100, where Wt is body weight at week *t* and Wi is the baseline body weight (set to 100%).

### Bulk Brain RNA-seq Sample Preparation, Data Processing and Analysis

Total RNA was extracted from frozen bulk brain tissue samples by Admera Health. RNA integrity was assessed using an Agilent TapeStation, and all samples had RNA Integrity Number (RIN) values > 8.5. RNA concentrations ranged from ∼275-606 ng/µL, with total yields of 11-24 µg. All samples were included for RNA-seq library preparation and paired-end sequencing. RNA-seq data processing and analysis were performed as previously described (50). Briefly, reads were aligned to the mouse reference genome GRCm39/mm10 using STAR (v2.7.11a). Transcript-level protein-coding counts were imported into R (v4.3.2) and analyzed with DESeq2 (v1.40.2). DEGs were defined as those with adjusted *p* value < 0.05 (Benjamini Hochberg correction) and |log2 fold change| ≥ 0.25. DEGs were visualized using volcano plots generated in ggplot2 (v3.4.4), displaying log2 fold change against -log10 p-value.

### Crude Synaptosome (P2 fraction) Preparation

Brain subcellular fractions were prepared by differential centrifugation as previously described (38, 51). A subset of mice (N=7/group) was deeply anesthetized using isoflurane and decapitated, their brains were immediately removed, then dissected in 2 hemispheres (cerebellum was discarded). A hemisphere from each animal was gently homogenized using a Teflon-glass homogenizer with 16-strokes at 800 rpm in 2 mL of Medium I (0.32 M sucrose, 5 mM HEPES pH 7.5, and 0.1 mM EDTA) containing protease and phosphatase inhibitors (ThermoFisher Scientific, 78446). Total brain homogenate was first centrifuged at 1,000 x g for 10 min at 4°C to give a pellet (P1) containing nuclear and cell debris. Supernatant (S1) was centrifuged at 12,000 x g for 20 min at 4°C to produce the crude synaptosome pellet (P2). Supernatant (S2) was discarded, P2 pellet was resuspended in 500 µL Medium I and sonicated on ice until completely dissolved, the resulting P2 fraction lysates were stored at -80°C until used for downstream analysis, including label-free quantitative mass spectrometry (LFQ‒MS), WB, and multiplexed Luminex immunoassays.

### LFQ‒MS Sample Preparation, Data Processing and Analysis

Total brain homogenate and P2 fraction lysates were centrifugated at 18,000 x g for 10 min at 4°C, supernatants were collected and protein concentration was determined by the BCA protein assay (ThermoFisher Scientific, 23225). Sample aliquots containing 50 µg of protein were prepared and digested according to the S-Trap micro spin columns protocol (PROTIFI, Document No. 20251212-08PRD-5021-BK-REV-B). Samples were adjusted to equal volumes with 8 M urea buffer (10mM TRIS, 100mM NaH2PO4, pH 8.5) and lysed in 5% SDS buffer, and then sonicated for 10 min, reduced with 10 mM dithiothreitol (Sigma, D5545) for 1 hour at RT, and alkylated with 20 mM iodoacetamide (Sigma, I6125) for 30 minutes at RT in the dark. Samples were acidified with phosphoric acid, and then diluted with binding buffer (100 mM triethyl-ammonium bicarbonate [TEAB] in 90% methanol pH 7.55). Next, samples were loaded onto S-Trap micro spin columns (PROTIFI, C02-micro-80) by centrifugation at 4,000 x g for 30 sec, and digested first with 1 µg lysyl endopeptidase (Wako, 127-06621) overnight at 37°C, followed by addition of 2 μg trypsin (ThermoFisher Scientific, 90058) with overnight incubation at 37°C. Peptides were sequentially eluted with 50 mM TEAB, followed by 0.2% formic acid, and finally 50% acetonitrile. The eluents were dried in a SpeedVac, dried peptides were resuspended in 0.1% formic acid and peptide concentration was determined by the quantitative fluorometric peptide assay (ThermoFisher Scientific, 23290). Peptides were subsequently analyzed by liquid chromatography–tandem mass spectrometry (LC‒MS/MS).

LC–MS/MS was performed on a Vanquish Neo UHPLC system coupled to an Orbitrap Exploris 480 mass spectrometer (Thermo Fisher Scientific) equipped with an EASY-Spray source, as previously described (48, 49). Briefly, 300 ng of peptides were loaded onto a C18 trap column (300 µm i.d. × 5 mm) and separated on an Aurora Frontier analytical column (75 µm i.d. × 60 cm, 1.7 µm C18; IonOpticks) at 300 nL/min. Mobile phases A and B consisted of 0.1% formic acid in water and 0.1% formic acid in 80% acetonitrile, respectively. The 135-min LC method used the following gradient: 1–4% B over 0–10 min, 4–25% B over 10–70 min, 25–65% B over 70–115 min, and 65–99% B over 115–116 min, followed by a hold at 99% B until 135 min. The column was re-equilibrated after each run.

MS data were acquired in positive-ion DIA mode with a spray voltage of 2.1 kV and an ion transfer tube temperature of 275 °C, without FAIMS. Each acquisition cycle comprised an MS1 scan over m/z 400–1000 at a resolution of 120,000, followed by 60 DIA MS2 scans at a resolution of 30,000. DIA window centers were spaced at 10 m/z intervals from m/z 405 to 995, with a 1 m/z overlap between adjacent windows (listed isolation ranges spanning 11 m/z). HCD fragmentation was performed at a normalized collision energy of 28%, and MS2 spectra were acquired over m/z 150–2000. Normalized AGC targets were 300% for MS1 and 1,000% for MS2, with automatic maximum injection times for both. Scan-to-scan internal mass calibration used user-defined lock masses at m/z 371.10124 and 445.12003.

Raw files were processed using DIA-NN v2.2.0 in library-free mode against the UniProt mouse database supplemented with contaminants. Searches used Trypsin/P specificity with up to two missed cleavages, fixed carbamidomethylation of cysteine, and common variable modifications. Results were filtered at 1% precursor and protein-group false discovery rates and quantified using QuantUMS without cross-run normalization.

Proteomic data from total brain homogenates and P2 fractions were processed in Perseus (http://www.perseus-framework.org/) as previously reported (38). Proteins were retained if at least 50% of values were present in one group, and intensities were log2-transformed. Missing values were imputed in Perseus by random sampling from a normal distribution. To correct sample loading differences, column-sum median normalization was performed in R (v4.3.2), where total protein intensity was calculated per sample, and each was scaled by the ratio of the global median column sum to its own sum. Group comparisons were assessed using one-way ANOVA and two-sample *t*-tests. Multiple testing correction was applied with the Benjamini Hochberg procedure (FDR < 0.05), with statistical significance defined at *p* < 0.05. DEPs were visualized using volcano plots generated in ggplot2 (v3.4.4), displaying log2 fold change against -log10 p-value.

Gene Ontology (GO) enrichment analysis was carried out using the R packages clusterProfiler (v3.21) and enrichR (v.3.4) packages, with the mouse genome annotation as the reference set and the experimentally identified proteome as the background. Enrichment was assessed across the three principal GO domains: Biological Process (BP), Molecular Function (MF), and Cellular Component (CC). Results were visualized in ggplot2 (v3.4.4), with terms ranked by -log10 adjusted p-value, and the most significantly enriched categories within each domain highlighted for downstream interpretation.

Gene Set Variation Analysis was performed using the GSVA R package (54) with a custom gene matrix transposed (GMT) file containing predefined gene sets. Pathway enrichment scores were calculated for individual samples using the z-score method. Pearson correlation analysis was performed to evaluate associations between pathway enrichment scores, with p-values adjusted for multiple testing using the Benjamini–Hochberg procedure. Correlation coefficients were visualized using heatmaps. To assess the robustness of pathway-level differences between experimental groups, permutation testing was performed using 1,000 random permutations of group labels. Empirical p-values were calculated as the proportion of permutations yielding absolute differences in mean pathway scores equal to or greater than the observed absolute difference.

### Western Blot (WB)

To confirm enrichment in synaptic elements, 8 µg of proteins from total brain homogenates and P2 fractions were resolved in a 4-12% Bis-Tris gel (Invitrogen, NW04125BOX) at 80V and transferred to nitrocellulose membranes using the iBLOT mini stack system (Invitrogen, IB23002). Membranes were incubated with Ponceau S staining solution (Cell Signaling, 59803) for 5 min to confirm equal loading of gels. After washing with 0.1% TBS-Tween 20, the blots were blocked with blocking buffer (ThermoFisher Scientific, 37543) for 1h at RT. Membranes were incubated with primary antibodies (Table 1A) for synaptic (SV2A, PSD95, RPH3A), mitochondrial (VDAC, TOMM20), cytosolic (α-tubulin), nuclear (histone H3), and astrocyte (GFAP) markers overnight at 4°C. Next day, after washing, membranes were incubated with corresponding secondary antibodies (Table 1B), washed and then imaged on the ChemiDoc Imaging System (Bio-Rad). The densitometric quantification of each band was performed using NIH ImageJ Fiji software (v2.16.0).

**Table 1.** List of primary and secondary antibodies used in this study.

| Target | Source | Cat # | Identifiers | Dilution | Application |
|---|---|---|---|---|---|
| PSD95 | Millipore | MAB1598 | RRID:AB_94278 | 1:500 | WB |
| SV2A | Abcam | ab32942 | RRID:AB_778192 | 1:1,000 | WB |
| RPH3A | Proteintech | 11396-1-AP | RRID:AB_2181145 | 1:1,000 | WB |
| VDAC | Cell signaling | 4661S | RRID:AB_10557420 | 1:1,000 | WB |
| TOMM20 | Abcam | ab186735 | RRID:AB_2889972 | 1:1,000 | WB |
| α-tubulin | Millipore | MAB1864 | RRID:AB_2210391 | 1:1,000 | WB |
| Histone H3 | Abcam | ab1791 | RRID:AB_302613 | 1:1,000 | WB |
| GFAP | Invitrogen | 13-0300 | RRID:AB_2532994 | 1:3,000<br>1:1,000 | WB<br>IF |
| Iba1 | Abcam | Ab178846 | RRID:AB_2636859 | 1:1,000 | IF |
| B) Secondary antibodies |  |  |  |  |  |
| Target | Source | Cat # | Identifiers | Dilution | Application |
| Goat anti-rabbit-Alexa 680 | Invitrogen | A-32734 | RRID:AB_2633283 | 1:10,000 | WB |
| Goat anti-rat-IRDye CW 800 | LI-COR | 926-32219 | RRID:AB_1850025 | 1:10,000 | WB |
| Goat anti-mouse-Alexa 800 | Invitrogen | A-32730 | RRID:AB_2633279 | 1:10,000 | WB |
| Donkey anti-rabbit Alexa 488 | Invitrogen | A-21206 | RRID:AB_2535792 | 1:500 | IF |
| Donkey anti-rat Alexa 647 | Invitrogen | A-78947 | RRID:AB_2910635 | 1:500 | IF |

WB indicates western blot; IF indicates immunofluorescence

### Quantification of Signaling Phospho-Protein Levels

Multiplexed Luminex immunoassays were used to measure phospho-proteins in the MAPK (Millipore, 48-660MAG) and mTOR pathways (Millipore, 48-612MAG) in total brain homogenates and P2 fractions as previously described (35, 36, 38). The MAPK 9-plex panel detected: pATF2 (Thr71), pErk (Thr185/Tyr187), pHSP27 (Ser78), pJNK (Thr183/Tyr185), pcJun (Ser73), pMEK1 (Ser222), pMSK1 (Ser212), p-p38 (Thr180/Tyr182), p-p53 (Ser15), pSTAT1 (Tyr701). The mTOR panel detected: pGSK3α (Ser21), pIGF1R (Tyr1135/Tyr1136), pIRS1 (Ser636), pAkt (Ser473), pmTOR (Ser2448), p70S6K (Thr412), pIR (Tyr1162/Tyr1163), pPTEN (Ser380), pGSK3β (Ser9), pTSC2 (Ser939), pRPS6 (Ser235/Ser236). A linear range analysis was conducted to identify the range of total protein loaded that corresponded to a linear signal readout from the Luminex instrument for each analyte (38, 55). Based on this analysis, 1.25 μg of total protein was loaded per sample for the MAPK panel, and 2.5 μg of protein for the mTOR panel. Luminex assays were read on an INTELLIFLEX system. Luminex data were z-scored and discriminant partial least squares regression analysis (D‒PLSR) was performed using the opls function in the ropls package in R (v.1.36.0). Standard deviation of latent variable one (LV1) and latent variable two (LV2) loadings for each protein was determined using a leave-one our cross validation approach. In this approach the D‒PLSR projection was recalculated 100 times with a random sample omitted each time. The output of D‒PLSR was rotated to best capture group-wise differences in LV1 and LV2.

### Histological Assessments

A subset of mice subjected to chronic SD and normal sleep controls (N=3/group) was deeply anesthetized using isoflurane and transcardially perfused with PBS followed by 4% paraformaldehyde. Brains were dissected in 2 hemispheres and post-fixed overnight at 4°C for histological assessments. A hemisphere was paraffin-embedded for Nissl staining and immunofluorescence, and the other hemisphere was processed for TEM.

### Nissl Staining

Paraffin-embedded hemispheres were sectioned using a Leica microtome, 10 μm-thick sagittal sections were deparaffinized at 60°C for 30 min and rehydrated, then stained with 1% cresyl violet (Abcam, ab246817) for 10 min, rinsed in distilled water, dehydrated, cleared in xylene, and cover slipped with Permount mounting medium (FisherScientific, SP15-100). Photomicrographs of the dorsal hippocampal CA1 and CA3 regions and neocortex consistently across all samples were acquired at 40x magnification using the Keyence BZ-X810 microscope. The number of pyramidal neurons showing normal morphology, distinct cytoplasmic and nuclear outlines, and visible nucleolus, was counted in a 100,687 µm^2^ area and then averaged to provide a single value for each animal (N=3 mice/group).

### Immunofluorescence (IF) microscopy

Following deparaffinization and rehydration, 5 μm-thick sagittal sections were boiled in sodium citrate buffer (pH 6) using a microwave during 10 min for antigen retrieval. After cooling, sections were permeabilized with 0.3% PBS-Triton X-100 for 10 min, and blocked with 5% Bovine Serum Albumin (BSA, Jackson ImmunoResearch, 001-000-162) for 2 h at RT. Then, sections were incubated overnight at 4°C with primary antibodies (Table 1A) for microglia (Iba1) and astrocyte (GFAP) markers in 1% BSA. Next day, after washing, sections were incubated with corresponding secondary antibodies (Table 1B) for 1 h at RT in the dark. After washing, sections were stained with DAPI (Vector Laboratories, H-1200) for 2 min, and mounted with antifade mounting medium (Invitrogen, P36970). IF images of the dorsal hippocampal CA1 and CA3 regions and neocortex were acquired at 20x magnification using the Keyence BZ-X810 microscope. A threshold level for each glial marker was established to detect differences between groups using NIH ImageJ Fiji software (v2.16.0). This threshold level was used for all images within a given marker. Percentage area covered indicate the area above the threshold level. The percentage areas were averaged to provide a single value for each animal (N=3 mice/group).

### TEM

First, fresh P2 fraction pellets were processed for ultrastructural analysis as previously described (38). Pellets were fixed in a mixture of 2.0% paraformaldehyde and 2.5% glutaraldehyde in 0.1 M cacodylate buffer (pH 7.4). The samples were next post-fixed with 1.0% osmium tetroxide (OsO4) and then stained en block in 2% aqueous uranyl acetate before being dehydrated through a series of graded ethanol concentrations up to 100% ethanol. Ethanol dehydration was followed by two changes in propylene oxide. Samples were then embedded in Eponate 12 resin and polymerized at 60°C for 48 h. Ultrathin sections were cut on a Leica EM UC6 ultramicrotome, stained with uranyl acetate and lead citrate, and imaged in a Hitachi HT7700 TEM, operated at 80KV. For quality assessment of crude synaptosomes, images were acquired at 5000x magnification.

Second, brain hemispheres (N=3 mice/group) were prepared for ultrastructural analysis of asymmetric synapses as previously described (38). After perfusion, hemispheres were quickly fixed using a mixture of 2.5% glutaraldehyde and 2% PFA in 0.1 M sodium cacodylate buffer (pH 7.4) for 1 h at RT and then kept overnight at 4°C. The next day, 150 µm-thick sagittal sections were cut on a vibratome, post-fixed for 1h on ice in the same fixative. The tissue was further post-fixed in 1% OsO4 and 0.8% potassium ferricyanide for 1 hour. Specimens were then en bloc stained with 2% aqueous uranyl acetate for 1 h, dehydrated in a graded series of ethanol up to 100%, substituted with propylene oxide, and embedded in EMbed 812 resin (Electron Microscopy Sciences). Sample blocks were polymerized in an oven at 60°C overnight. Ultrathin sections (60 nm) were cut using a Leica UC7ultramicrotome and post-stained with 2% uranyl acetate and lead citrate. The ultrathin sections were examined with an FEI Tecnai transmission electron microscope at an 80 kV accelerating voltage, and digital images were acquired with an AMT NanoSprint15 MK2 camera (Advanced Microscopy Techniques). For the image analysis, the asymmetric synapse was defined as an electron-dense post-synaptic density area juxtaposed to a presynaptic terminal. Images used for analysis were acquired from hippocampal CA1 region at 6800x magnification, and the analysis was performed while blind to the experimental conditions. Morphological features including presynaptic bouton size (area) and length of postsynaptic density (PSD) were manually delineated and measured using NIH ImageJ Fiji software (v2.16.0), while the number of synaptic vesicles was manually counted and then divided by the presynaptic bouton area to estimate synaptic vesicle density.

### ELISA

Prior to euthanasia by decapitation or transcardiac perfusion, mice were deeply anesthetized with isoflurane. Blood was collected in EDTA tubes and then centrifuged at 1,000 x g for 10 min at 4°C to obtain plasma. Plasma samples were diluted [1:40] and [1:25] in assay buffer respectively to measure Corticosterone and NfL levels using commercially available ELISA kits (Enzo, ADI-900-097; Umon Diagnostics, 20-8002) following the manufacturer’s protocols.

### Statistical analysis and data visualization

RNAseq and proteomic data analyses were conducted in R. Statistical analyses were performed using GraphPad Prism (v.10.2.3). Data are reported as mean ± standard error of the mean (SEM), with significance set at *p* < 0.05. Graphs were prepared using GraphPad Prism (v.10.2.3), schematics were created using BioRender.com, and figures were edited using Adobe Illustrator 2024 (v.28). Specific statistical considerations are provided in the corresponding figure legends.

## RESULTS

### Global effects of the chronic SD paradigm in male WT mice

By using previously validated sleep fragmentation chambers (39, 45–49), we investigated the cumulative effects of chronic SD on brain transcriptomic signatures and compartment-specific proteomic profiles to identify genes, proteins, and signaling pathways that might confer AD risk. In the chronic SD paradigm, 6.5 month-old male WT mice were group-housed in automated sleep fragmentation chambers equipped with a bar set to move every 30 sec for 6 h/day, from ZT3 to ZT9, over 6 consecutive weeks **(Fig. 1A)**. As expected (39, 49), mice in the SD group exhibited persistent lower body weight compared to mice in the undisrupted sleep control group, while no alterations in plasma corticosterone levels were observed at the time of euthanasia **(Fig. 1B)**. These results suggest that the body weight changes observed in SD mice might be attributable to increased locomotor activity, which is intrinsic to this paradigm, or to direct SD effects, rather than to a stress response. No animals died or exhibited overt health decline throughout the experiment.

**Figure 1.**
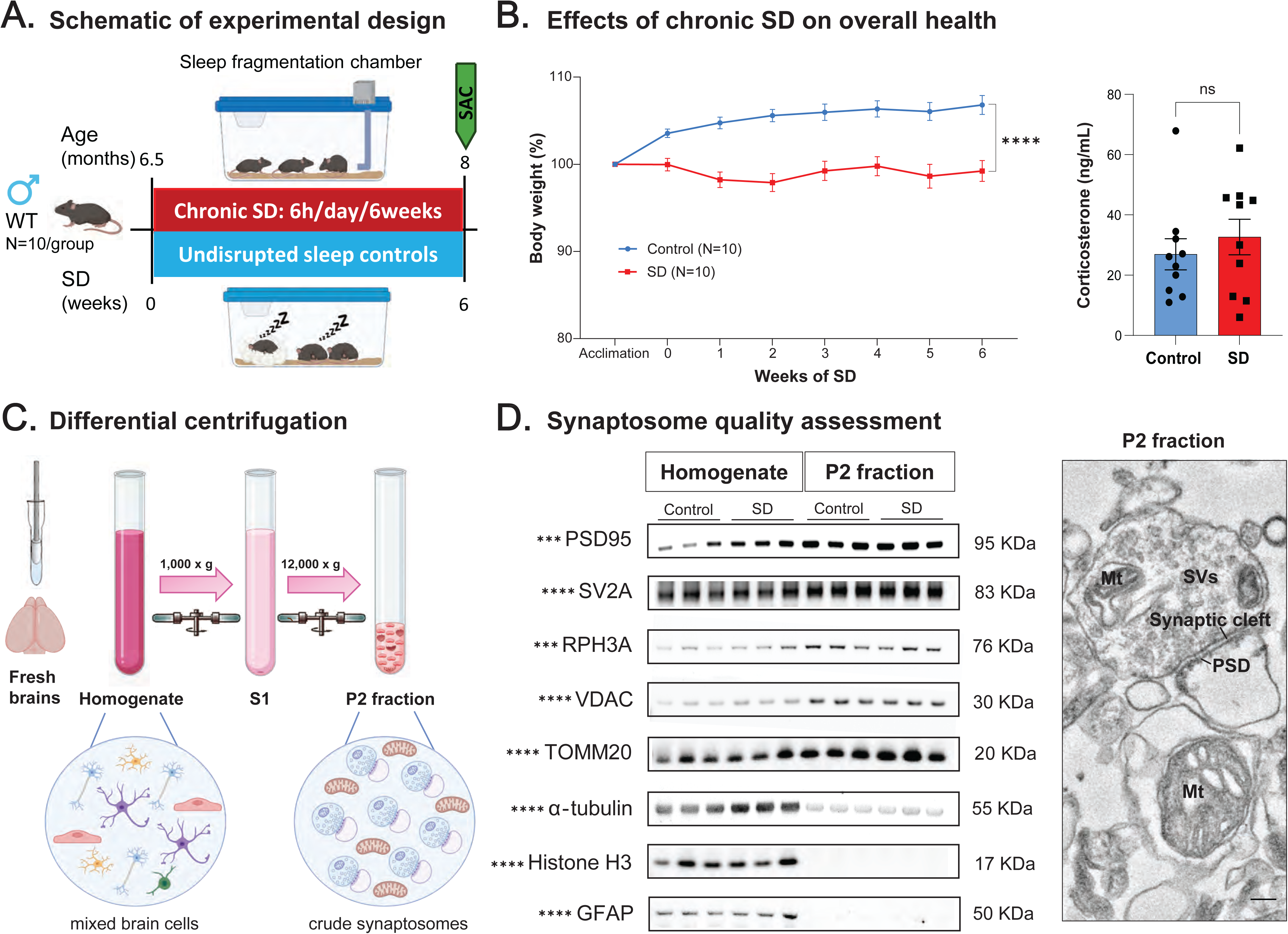
Overview of the study and experimental design. **(A)** Chronic sleep disruption (SD) paradigm. **(B)** Chronic SD significantly reduced body weight gain, but did not alter plasma levels of corticosterone. Data are expressed as mean ± SEM. Left: Two-way repeated measures ANOVA [F(3.093, 55.68)=11.90, \*\*\*\**p*<0.0001]. Right: Unpaired t-test [t=0.7313, df=18, p=0.474], ns = no significant. **(C)** Differential centrifugation method to prepare crude synaptosomes (P2 fractions). **(D**, left) Representative blots showing enrichment of synaptic (PSD95, SV2A, RPH3A) and mitochondrial (VDAC, TOMM20) proteins in P2 fractions compared to brain homogenates, while depletion of cytosolic (α-tubulin), nuclear (histone H3), and astrocytic (GFAP) markers. Mean fold change values (P2/H) were calculated (N=14) and analyzed by one sample t-test [PSD95: t=4.529, df=13; SV2A: t=6.668, df=13; RPH3A: t=5.427, df=13; VDAC: t=7.383, df=13; TOMM20: t=10.03, df=13; a-Tubulin: t=15.20, df=13; H3: t=200.7, df=13; GFAP: t=37.85, df=13; \*\*\**p*<0.001, \*\*\*\**p*<0.0001]. **(D**, right) Representative electron micrographs of P2 fractions confirmed the presence of synaptic vesicles (SVs), mitochondria (Mt), and post-synaptic density (PSD). Scale bar: 200 nm. Also see Supplemental Table S1 for related analyses and datasets. Schematics in Figs. 1A and C were created using BioRender.com.

We next examined the effects of the chronic SD paradigm on neuronal density and glial responses in the brain (**Fig. 2, Supplemental Table S1**). Mice in the SD group did not show alterations in the number of pyramidal neurons in hippocampal CA1 and CA3 regions nor the neocortex (**Fig. 2A-B**). Also, no changes were observed in glial markers. SD mice and normal sleep control mice displayed similar percentages area covered by Iba1+ myeloid cells and GFAP+ astrocytes in the hippocampal CA1 and CA3 regions and in the neocortex (**Fig. 2A-C**). No changes were observed in plasma NfL levels of SD mice at the time of euthanasia (**Fig. 2D**). These results indicate that the chronic SD paradigm used in this study did not induce pronounced neuropathological changes (i.e., neurodegeneration, neuroinflammation or axonal injury) in male WT mice.

**Figure 2.**
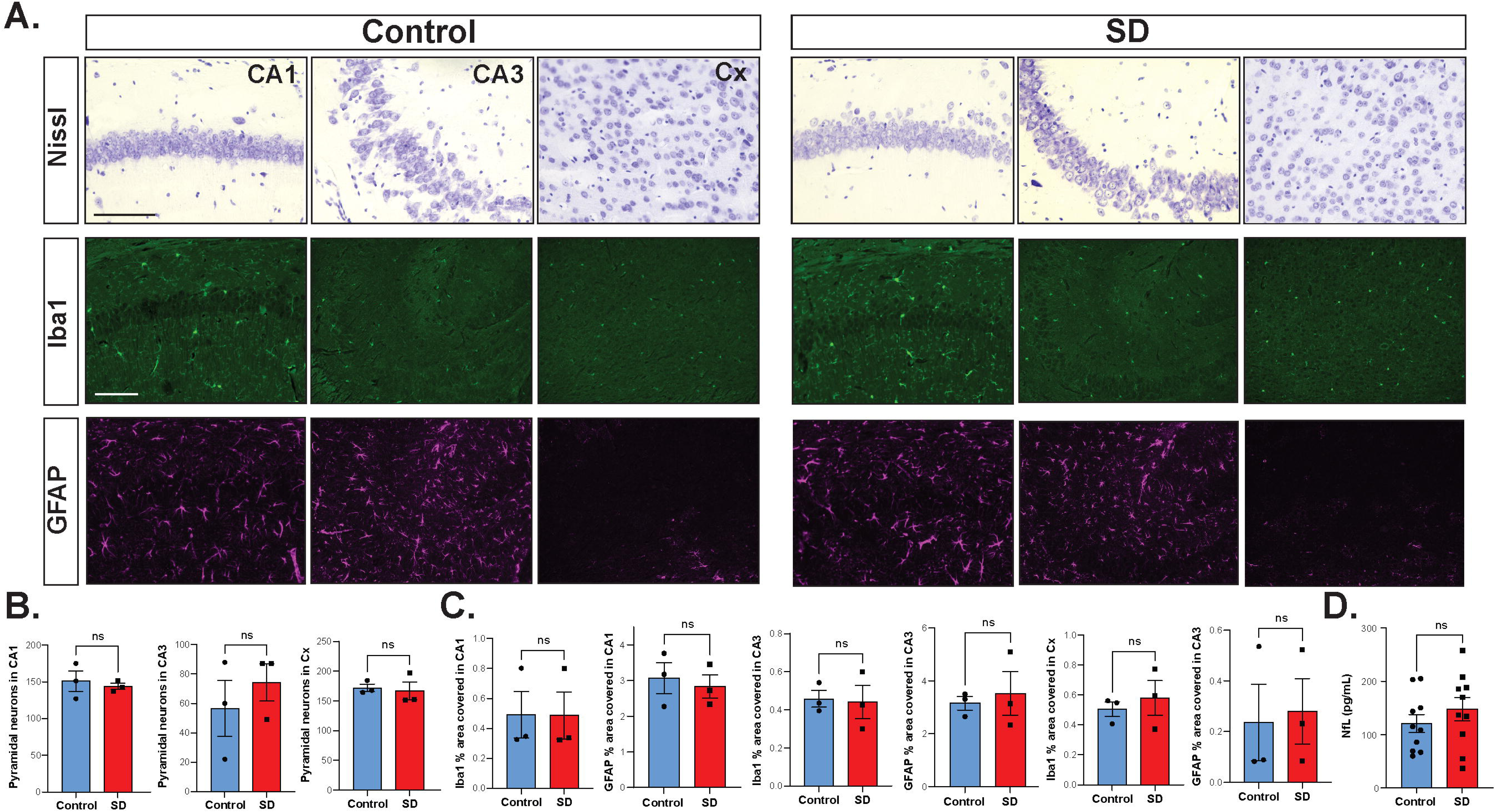
Effects of chronic SD on neuronal density and glial marker responses in the brain. **(A**, top panel) Representative photomicrographs of Nissl-stained sagittal sections showing hippocampal CA1 and CA3 regions and neocortex (Cx) from mice subjected to chronic SD, or from undisrupted sleep controls. **(A**, middle panel) Representative immunofluorescence images showing the distribution of Iba1^+^ cells (microglia in green), and **(A**, bottom panel) GFAP^+^ cells (astrocytes in magenta) in hippocampal CA1 and CA3 regions and neocortex from mice subjected to chronic SD, or from undisrupted sleep controls. Scale bars: 100 μm. **(B)** Number of pyramidal neurons in hippocampal CA1 and CA3 regions and neocortex. Chronic SD did not modify neuronal density in these brain regions. Data are expressed as mean ± SEM. Unpaired t-test [CA1: t=0.5058, df=4, p=0.6396; CA3: t=0.7701, df=4, p=0.4842; Cx: t=0.3311, df=4, p=0.7572], ns = no significant. **(C)** Percentage area covered by Iba1 and GFAP in hippocampal CA1 and CA3 regions and neocortex. Chronic SD did not alter glial markers in these brain regions. Data are expressed as mean ± SEM. Unpaired t-test [Iba1 CA1 % area: t=0.01960, df=4, p=0.9853; GFAP CA1 % area: t=0.4214, df=4, p=0.6951; Iba1 CA3 % area: t=0.1644, df=4, p=0.8774; GFAP CA3 % area: t=0.4271, df=4, p=0.6913; Iba1 Cx % area: t=0.5924, df=4, p=0.5855; GFAP Cx % area: t=0.2241, df=4, p=0.8337], ns = no significant. **(D)** Chronic SD did not alter plasma levels of neurofilament light chain (NfL). Data are expressed as mean ± SEM. Unpaired t-test [t=0.9613, df=18, p=0.3491], ns = no significant. Also see Supplemental Table S1 for related analyses and datasets.

### Chronic SD triggers transcriptional cellular stress and the unfolded protein response

Acute SD is known to alter gene expression, molecular signaling, and structure in the rodent brain. For instance, acute SD strongly represses gene transcription, downregulating thousands of genes in the rodent brain (32, 56, 57). However, the cumulative effects of chronic SD remain poorly defined. We performed bulk brain RNAseq to identify transcriptomic effects of chronic SD imposed by this paradigm in male mice. Bulk transcriptomic analysis of 17,574 transcripts revealed minimal gene changes in the brain after chronic SD at the BH FDR<0.05 significance level. These included 4 DEGs, 2 upregulated genes (*Hspa1a, Hspa1b*) and 2 downregulated genes (*Nrp1, Tnxb*) (**Fig. 3A**). These transcriptional changes suggest an increased expression of chaperones/heat shock proteins in response to cellular stress and activation of the unfolded protein response.

**Figure 3.**
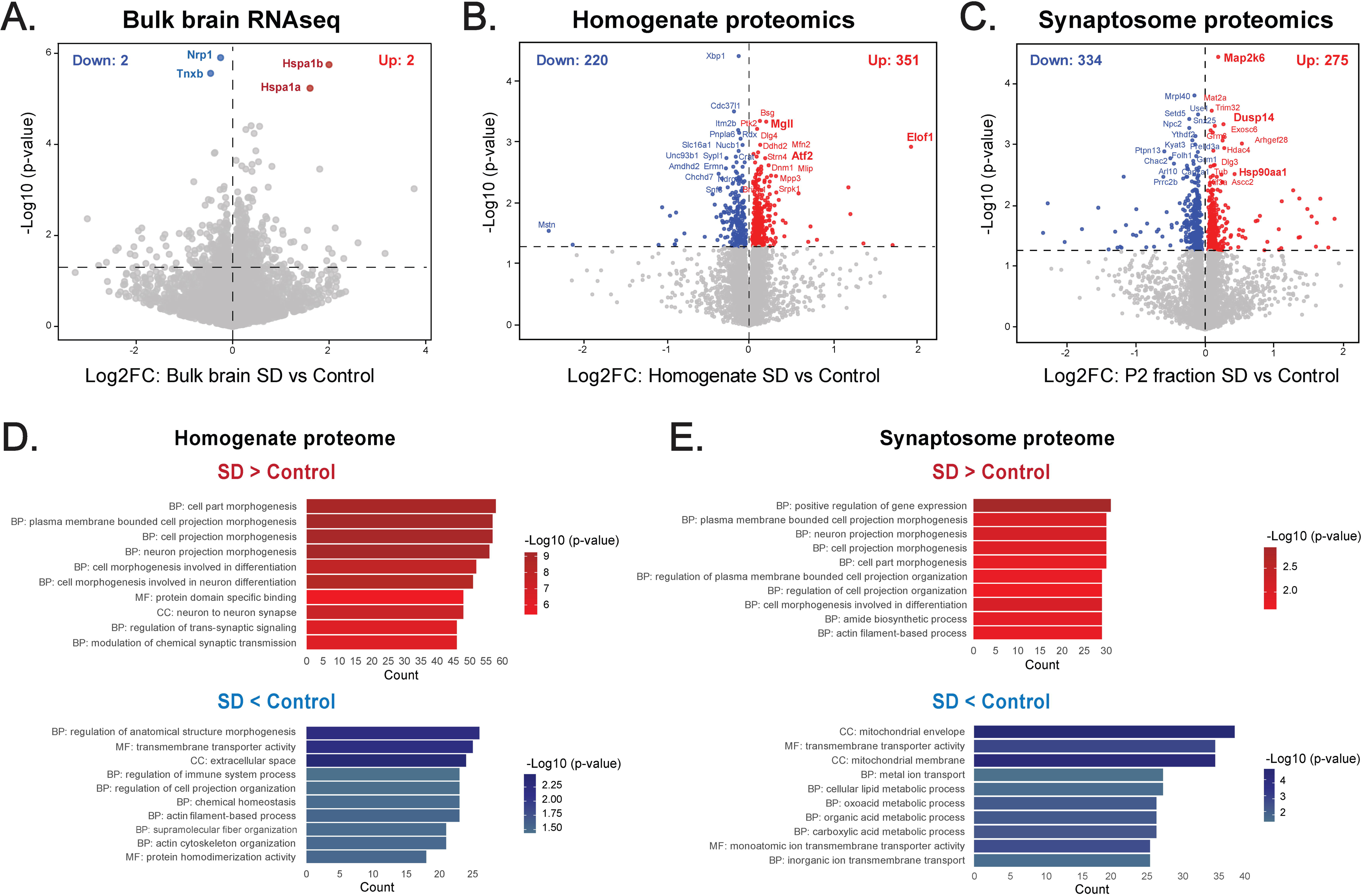
Brain transcriptomic and compartment-specific proteomic effects of chronic SD. **(A)** Volcano plot showing differentially-expressed genes (DEGs) in bulk brain samples from mice subjected to chronic SD compared to undisrupted sleep controls (N=7/group). Volcano plots showing differentially-enriched proteins (DEPs) in **(B)** whole-brain (homogenate) samples and on **(C)** synaptosome (P2 fraction) samples from mice subjected to chronic SD compared to undisrupted sleep controls (N=7/group). **(D)** Gene Ontology (GO) enrichment analyses of homogenate (SD vs Control) proteomes and **(E)** synaptosome (SD vs Control) proteomes. Selected GO terms from each of the three GO groups (biological process [BP], cellular component [CC], and molecular function [MF]) are shown. Also see Supplemental Table S1 for related analyses and datasets.

### Chronic SD induces compartment-specific proteomic effects

To investigate compartment-specific proteomic effects of chronic SD, we employed a subcellular fractionation method via differential centrifugation to prepare crude synaptosomes (synapse-enriched P2 fractions) (**Fig. 1C**). WB analysis confirmed that P2 fractions were significantly enriched in synaptic (PSD95, SV2A, RPH3A) and mitochondrial (VDAC, TOMM20) proteins, while depleted in cytosolic (α-tubulin), nuclear (histone H3), and glial (GFAP) markers (**Fig. 1D, *left panel***). TEM further confirmed that the P2 fractions were enriched in pinched-off pre-synaptic terminals containing synaptic vesicles, post-synaptic elements such as the PSD, and mitochondria (**Fig. 1D, *right panel***).

Following biochemical and ultrastructural validation of P2 fractions, we then performed LC‒MS/MS on P2 fractions and whole-brain homogenates and identified 6,985 proteins. Principal-component analysis (PCA) of the P2 fraction and homogenate proteomes revealed separated clustering between compartments along the first PC, and additional separation among experimental groups (**Supplemental Table S1**).

Chronic SD induced 571 DEPs (351 increased and 220 decreased) in the homogenate proteome (**Fig. 3B**), and 609 DEPs (275 increased and 334 decreased) in the P2 fraction proteome (**Fig. 3C**), of which only 53 DEPs overlapped (**Fig. 4A**). Among the overlapping DEPs, Hspa1b levels were elevated, consistent with its upregulated gene expression observed in the bulk transcriptomic analysis (**Fig. 3A**). This concordance in Hspa1b upregulation supports that chronic SD induces the activation of the unfolded protein response in the brain.

**Figure 4.**
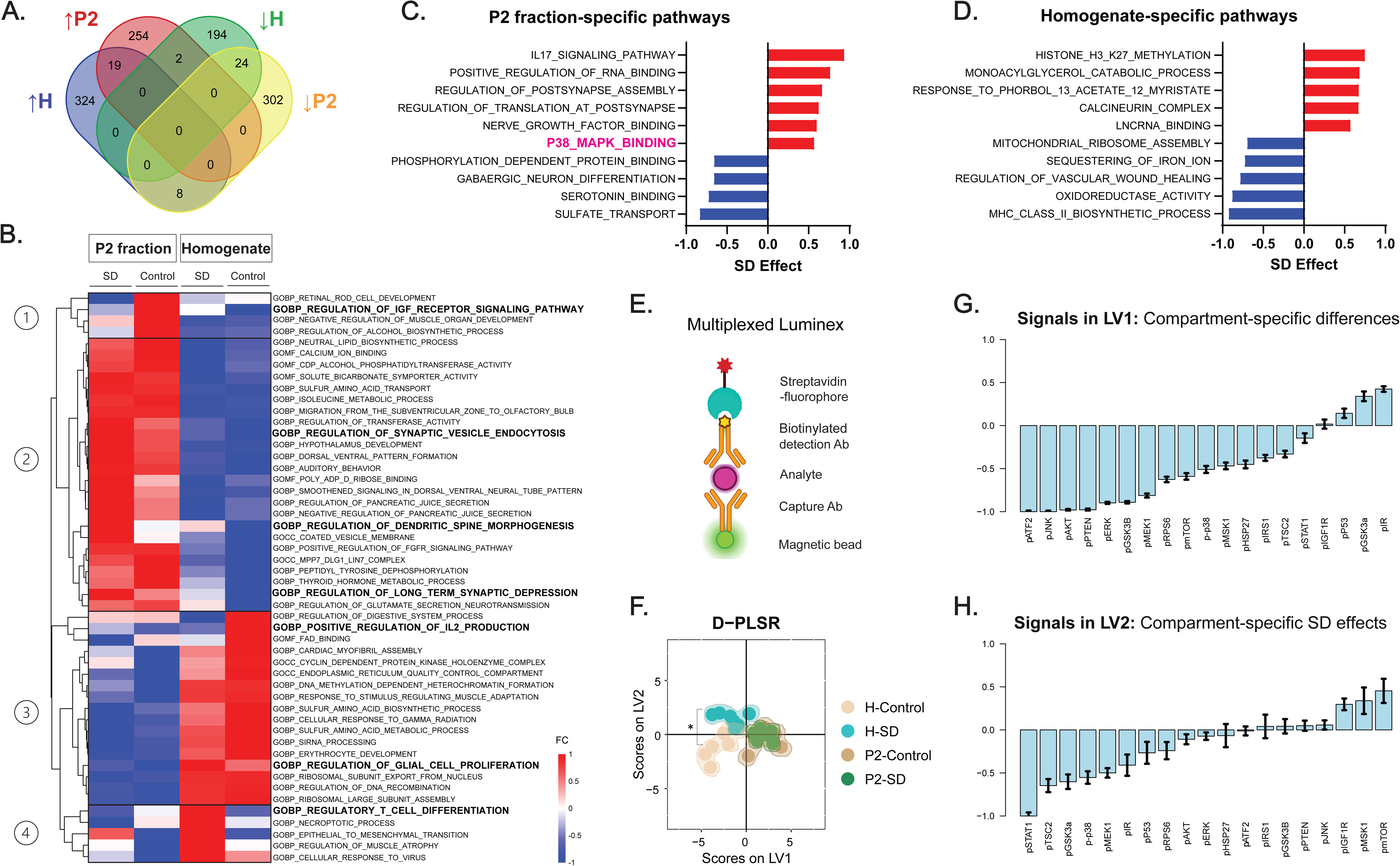
Compartment-specific proteomic and signaling effects of chronic SD. **(A)** Venn diagram of the unique and shared differentially-enriched proteins (DEPs) identified from brain homogenates and P2 fractions. **(B)** Gene set variation analysis (GSVA) reveals differential effects of chronic SD on pathway enrichment in homogenates and P2 fractions. Top 10 GSVA pathways dysregulated by chronic SD specifically in **(C)** P2 fraction and **(D)** homogenate proteomes. **(E)** Cartoon representation of multiplexed Luminex assay to measure the levels of phospho-proteins belonging to MAPK and Akt/mTOR signaling pathways from homogenates and P2 fractions. **(F)** D-PLSR analysis was used to cluster the samples based on treatment and sample type. Wilcoxon rank sum test [H-Control vs H-SD: W=1, p=0.0012; P2-Control vs P2-SD: W=23, p=0.9015] Individual phosphoprotein analyte loadings on **(G)** LV1 and **(H)** LV2: Positive values indicate the analyte is associated with positive scores on a given LV and vice versa for negative loadings. Error bars indicate standard deviation of loadings from leave-one out cross validation. Also see Supplemental Table S1 for related analyses and datasets. Schematic in Fig. 4E was created using BioRender.com.

To decipher the functional implications of these protein alterations, we then conducted gene ontology (GO) enrichment analysis and gene set variation analysis (GSVA) from the upregulated and downregulated proteins in both compartments. On the one hand, GO analysis in the homogenate proteome indicated that chronic SD increased levels of proteins involved in neuron morphogenesis (Tanc2, Shank1, Syngap1, Ephb6, Mgll, Src, App, Cdh2), synaptic signaling (Mapk1, Dlg4, Ptprs, Gnai1, Dnm1, Grip1) and chemical synaptic transmission (Nrxn2, Nr3c1, Nptn, Pla2g6, Epha4, Synpr, Shisa7, Mpp2). Conversely, chronic SD reduced levels of proteins involved in transmembrane transporter activity (S100a1), regulation of immune system process (Lamp1, Cd81, Bax, Notch2, Il33), actin cytoskeleton reorganization (Epb41l2, Pls1, Anln, Rhoq) (**Fig. 3D**). On the other hand, GO analysis in the P2 fraction proteome indicated that chronic SD increased levels of proteins involved in positive regulation of gene expression (Hspa1b, Hsp90aa1, Trim32, Exosc6, Gsk3b, Eif3e), neuron projection morphogenesis (Sema4c, Ncam1, Plxnb3, Rac1, Cdh4, Flot1), and differentiation (Hsp90aa1, Rab3a, Syngap1, Pak3, Il1rapl1, Mark2). Conversely, chronic SD reduced levels of proteins involved in mitochondrial envelope (Mrpl40, Pgrmc1, Vdac1, Nelfb, Slc25a21, Coa7, Abca8b), cellular lipid metabolic process (Cln8, Ucp3, Lmf1, Asah2, Gpld1, Ptgr2), and carboxylic acid metabolic process (Scd2, Ucp3, Ptgr2, Acox3, Spr, Kyat3) (**Fig. 3E**). Conserved effects of chronic SD included increased chaperones (Hspa1b) and excitatory synaptic proteins (Dlg3, Syngap1, Dkk3, Mpp3), but reduced inhibitory GABAergic receptors (Gabrg1), ion channels (Kcnc3, Slc6a5), and neuronal structural proteins (Map4, Nefl) at both whole-brain and synaptosome proteomic levels (**Supplemental Table S1**).

Importantly, Venn diagram of DEPs shows minimal overlap of chronic SD effects across the two compartments (**Fig. 4A**), indicating that the effects of chronic SD on homogenate and P2 fraction proteomes are molecularly distinct. Similarly, GSVA analysis of the homogenate and P2 fraction proteomes revealed that chronic SD differentially altered pathway enrichment in the synaptic compartment (clusters 1 and 2), compared to its effects in the whole-brain proteome (clusters 3 and 4) (**Fig. 4B**). For instance, chronic SD increased key cellular pathways in the homogenate, including regulatory T-cell differentiation, necroptotic process, and regulation of glial cell proliferation, whereas decreased positive regulation of IL-2 production (**Fig. 4B**). In contrast, in P2 fractions, chronic SD increased proteins involved in regulation of synaptic vesicle endocytosis, regulation of dendritic spine morphogenesis, and regulation of long-term synaptic depression, but reduced proteins involved in regulation of IGF receptor signaling pathway (**Fig. 4B**). Interestingly, top GSVA pathways illustrate compartment-specific dysregulation by SD in P2 fraction and homogenate proteomes. Specific to the P2 compartment, chronic SD increased IL-17 pro-inflammatory signaling, regulation of local translation, and p38 MAPK binding, while reducing phosphorylation dependent protein binding and GABAergic neuron differentiation (**Fig. 4C**). In contrast, specific to the homogenate, chronic SD increased histone methylation and monoacylglycerol catabolic process, while reducing mitochondrial ribosome assembly and oxidoreductase activity at the whole brain level (**Fig. 4D**). These findings suggest that chronic SD-induced protein alterations might reflect a compensatory signature, where synaptic-associated pathways were particularly prominent in the P2 fraction proteome, while immune/inflammatory, glial responses and neuronal function pathways were mostly enriched in the homogenate proteome.

### Chronic SD negatively regulated the MAPK signaling pathway

Besides its impacts on transcription and translation, chronic SD has been reported to alter signaling through post-translational modifications, such as protein phosphorylation (28, 32, 58). The mitogen-activated protein kinases (MAPK) and Akt/mTOR signaling pathways play an important role in how disrupted sleep impairs synaptic function and plasticity (59, 60). Upon activation at the synapse, MAPK and mTOR phosphorylate downstream kinases, which regulate key transcriptional factors inducing local translation of target genes involved in synaptic plasticity, including scaffolding proteins and neurotransmitter receptors (61, 62). To examine the impact of chronic SD on these signaling pathways, we used multiplexed Luminex immunoassays (**Fig. 4E**) for measuring phospho-protein levels in homogenates and P2 fractions. At control conditions (undisrupted sleep controls), D‒PLSR analysis revealed that MAPK and Akt/mTOR pathway activation was lower in the synaptic compartment (exemplified by a negative signal of LV1 for pATF2, pJNK, pAKT, pERK, pmTOR, p-p38 in the P2 fractions) as compared to the whole-brain level (**Fig. 4F-G**). Notably, we observed that the effect of chronic SD on these signaling pathways was most apparent at the whole-brain level as opposed to the synaptic compartment, highlighting that chronic SD suppressed MAPK pathway activation in the homogenates (exemplified by a negative signal of LV2 for pSTAT1, pMEK1, p-p38, pERK) compared to the P2 fractions, while Akt/mTOR signaling was not impacted (**Fig. 4F-H**, **Supplemental Table S1**).

### Chronic SD subtly alters the ultrastructure of vulnerable hippocampal CA1 synapses

Next, we investigated whether chronic SD-induced proteomic and signaling alterations might also affect synaptic ultrastructure. Using TEM, we examined asymmetric synapses in the hippocampal CA1 region, a brain area particularly vulnerable to sleep deprivation (63, 64) (**Fig. 5A**). When comparing individual synapses between SD and Control groups, the cumulative distribution curves revealed that presynapse size (**Fig. 5B**) and synaptic vesicle density (**Fig. 5C**) were significantly decreased in SD mice, while the PSD length did not change (**Fig. 5D**), as compared to undisrupted sleep control mice. These results showing decreased presynapse size and synaptic vesicle density, suggest that chronic SD modifies synaptic transmission and plasticity, consistent with alterations of important synaptic proteins in the P2 fraction (e.g., reduced levels of Grm1, Unc13c, Sema4c, Vdac1, Ncam1, Kcnc3, Itga3) and the homogenate (e.g., reduced levels of Cdk5r1, Pls3, Adora2a, Gabrg1, Adam10) proteomes from SD mice (**Supplemental Table S1**).

**Figure 5.**
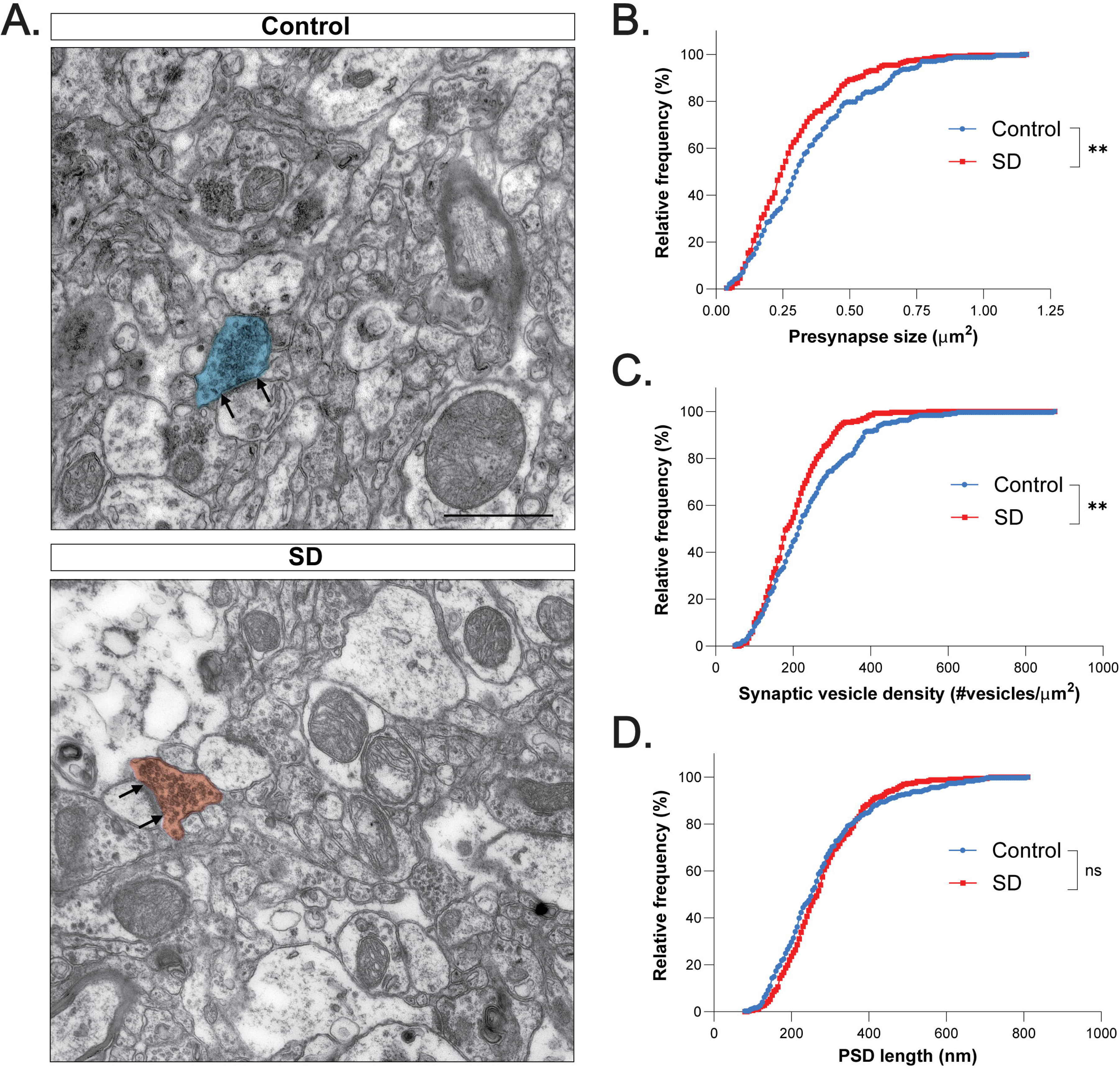
Effects of chronic SD on the synaptic ultrastructure. **(A)** Representative electron micrographs showing asymmetric synapses in hippocampal CA1 region from mice subjected to chronic SD compared to normal sleep controls. Scale bar: 1 µm. Cumulative distribution of **(B)** presynapse size, **(C)** synaptic vesicle density and **(D)** PSD length. Kolmogorov–Smirnov (K-S) test [Presynapse size: D=0.1677, p=0.0019; Synaptic vesicle density: D=0.1598, p=0.0036; PSD length: D=0.09485, p=0.1265], \*\**p*<0.01, ns = no significant. Control N=236 and SD N=261 asymmetric synapses from three WT mice for each group. Also see Supplemental Table S1 for related analyses and datasets.

## DISCUSSION

Given its high prevalence in modern society, SD represents an important risk factor for the development and progression of AD (8, 14–23). However, the molecular mechanisms underlying this association are not completely understood, especially at synapses, which are particularly vulnerable to sleep loss. Although previous studies have shown that acute SD alters synaptic molecular composition, ultrastructure, and function, critical gaps remain in our understanding of how chronic SD triggers pathological changes in the brain and its vulnerable synapses driving disease risk and neurodegeneration. Therefore, deciphering the molecular alterations induced by chronic SD is a crucial step towards identifying potential targets for therapeutic intervention.

Here, we used bulk brain transcriptomics, bulk proteomics and synaptosome-enriched proteomics, multiplexed Luminex immunoassays of signaling pathways and structural imaging of synapses, to characterize the cumulative effects of chronic SD on genes, proteins, and signaling pathways at the whole-brain level and the synaptic compartment. Our findings demonstrate that chronic SD triggered a minimal transcriptional response (only 4 DEGs) in the brain characterized by increased gene expression of chaperones/heat shock proteins associated with cellular stress and the unfolded protein response. In contrast with the minimal transcriptional response, the effects of chronic SD on the brain and the synaptic compartment proteomes were more extensive, with 571 DEPs identified in homogenate and 609 DEPs in the P2 fraction. Remarkably, these proteomic alterations were compartment-specific, highlighting upregulated pathways such as synaptic vesicle endocytosis, regulation of dendritic spine morphogenesis, and regulation of long-term synaptic depression that might indicate compensatory mechanisms in the synaptic compartment proteome, contrasting with upregulated immune/inflammatory pathways, glial responses and neuronal function observed in the whole-brain proteome. Next, we complemented proteomics data using multiplexed Luminex to measure phospho-proteins of signaling pathways relevant for synaptic function and plasticity, and found that chronic SD suppressed MAPK signaling at the whole brain level, but no significant effect was observed in the synaptic compartment. Consistent with the synaptic protein alterations observed in proteomics data, chronic SD subtly altered the ultrastructure of vulnerable hippocampal CA1 synapses, reducing presynapse size and synaptic vesicle density that might contribute to impaired synaptic transmission and plasticity. Collectively, these results identify molecular candidates (e.g., Hspa1b, Map2k6, and Atf2) and the p38 MAPK pathway as potential therapeutic targets. It is worth noting that increased levels of Map2k6 (65) and p38 MAPK activation (66) have been reported to occur in hippocampal vulnerable neurons of AD patients. However, future mechanistic studies of chronic SD in AD-relevant mouse models are needed to determine whether targeting these factors can attenuate synaptic dysfunction at the early stages of AD.

### Impact of chronic SD on brain gene expression responses

Numerous preclinical studies have demonstrated that SD induces changes in gene expression, protein translation and signaling (28, 32, 58). Particularly, in the context of acute SD, transcriptomic studies have reported altered expression of thousands of genes in the rodent brain. For instance, Ford et al. (57) found that acute SD strongly represses transcription at the gene and transcript level in the frontal cortex, preferentially affecting glutamatergic neurons (1492 DEGs). Moreover, studies using spatial transcriptomics (67), single-nuclear RNAseq (57) and combining cell-type specific transcriptomics with nuclear proteomics (68) have reported unique cell-type and region-specific transcriptional signatures in response to acute SD. Further, Gaine et al. (69) identified 1,146 genes that were significantly dysregulated following acute SD (5 h via gentle handling), with 507 genes upregulated (associated with RNA splicing and the nucleus) and 639 genes downregulated (associated with cell adhesion, dendritic localization, the synapse, and postsynaptic membrane). Importantly, most DEGs normalized after 3 h recovery sleep. Similarly, Popescu et al. (56) found that acute SD-induced gene expression changes in the cortex peaked at 5-6 h of SD, including genes involved in cellular redox homeostasis, DNA damage/repair, and chromatin regulation. Also in their study, most DEGs normalized after 2 hours of recovery sleep, except genes associated with cellular stress.

In contrast with acute SD findings, early studies from Cirelli et al. (70), using microarray labeling and hybridization in rats subjected to the disk-over-water method for 7 days, reported that chronic SD increased 75 transcripts coding for immune and stress response proteins, and decreased 16 transcripts coding for proteins involved in connective tissue integrity and energy metabolism. Furthermore, Ba et al. (71), using a 12 h/day sleep fragmentation paradigm for 45 days, found that chronic SD induced 98 DEGs in the hippocampus, which were mainly involved in neuroinflammation, immune system dysfunction, metabolic and molecular transcription dysregulation. These gene expression changes were accompanied of AD-like pathological changes including tauopathy and gliosis. In this study, using a 6 h/day sleep fragmentation paradigm for 42 days (6 weeks), we did not observe significant pathological changes in the hippocampus nor the neocortex, and bulk brain transcriptomic analysis identified only 4 DEGs after chronic SD that predominantly encoded chaperone/heat-shock proteins associated with cellular stress and the unfolded protein response. In line with Gaine et al. (69) and Popescu et al. (56), the minimal transcriptional response observed here might be consequence of DEGs normalization reported to occur after an extended duration of recovery sleep (18 h before brain tissue collection).

Among DEGs, we found upregulation of Hspa1a and Hspa1b which encode HSP70. Interestingly, recent evidence supports the therapeutic potential of targeting HSP70 in the context of chronic SD. For example, Kang et al. (72) engineered exosomes to deliver HSP70 mRNA specifically to the brain of mice subjected to the modified multiple-platform water paradigm for 20 h/day, over 7 consecutive days, and demonstrated that HSP70 has neuroprotective effects by reversing SD-induced cognitive impairments and reducing inflammation in the hippocampus.

### Impact of chronic SD on brain and synaptosome proteomes

Omics studies of sleep can reveal critical targets and molecular pathways impacted by SD in health and disease (73, 74). In fact, it is well documented that SD alter brain protein synthesis, folding, degradation, and post-translational modifications (28, 32). Synaptosomes are efficient tools to identify disease-associated molecular mechanisms occurring at the synapses (75, 76). Recent proteomic studies have used synaptosomes to characterize the effects of SD on synaptic protein composition. For instance, Simor et al. (77) reported that acute SD (8 h via gentle handling) induced short- and long-term proteomic changes in cortical and thalamic synapses, identifying 142 DEPs, of which 125 were unique to cortical synapses, including proteins involved in synaptic function and plasticity (e.g., synaptic vesicle recycling, dendritic outgrowth and synaptogenesis). Similarly, Gulyássy et al. (30) showed that acute SD (8 h via gentle handling) induced 78 DEPs (56 upregulated and 22 downregulated) in cortical synapses, dysregulated proteins were involved in signal transduction and regulation, synaptic assembly and neurotransmission, lipid metabolism, cytoskeleton organization, protein synthesis, degradation and folding. Interestingly, after 16 h of recovery sleep, 39 DEPs persisted (15 upregulated and 24 downregulated). In addition, Noya et al. (78) quantified 4,063 synaptic proteins in forebrain synaptosomes and found that 474 (11.7%) exhibited 24-hour rhythmic abundance, including proteins implicated in synaptic plasticity and neuronal metabolism. Strikingly, acute SD (4 h via gentle handling) dampened rhythmicity cycles in ∼98% of those synaptic proteins. Recently, Gay et al. (27) characterized the effects of acute SD (4 h or 6 h via gentle handling) on the forebrain synapse proteome and phosphoproteome. Proteomic analysis identified 326 altered synaptic proteins, with 5.9% regulated by 4 h-SD, while 772 altered proteins, with 11.7% regulated by 6-h SD. Contrary to this, the phosphoproteomic analysis showed that 4 h-SD induced a modest response of 491 (2.6%) phosphopeptides in comparison with 6 h-SD response of 1,845 regulated phosphopeptides (11.4%). These findings demonstrate that acute SD exerts distinct molecular consequences on the forebrain synapses depending on its duration. It is worth noting that the observed discrepancies in the number of identified DEPs among studies might result from differences in synaptosome preparation protocols (e.g., ultracentrifugation versus Percoll-gradient fractionation) and sample purity (e.g., crude versus purified synaptosomal fractions) (79).

A key finding of this study is that chronic SD induced compartment-specific proteomic alterations, with a different impact on the synaptic proteome, showing increased proteins involved in synaptic vesicle endocytosis (Rab3a, Numb, Hpca, Grb2, Gria2), regulation of dendritic spine morphogenesis (Itpka, Pak3, Gsk3b, Arc), and regulation of long-term synaptic depression (Shank2, Fmr1, Actr2), compared to the whole-brain proteome, including increased proteins involved in immune/inflammatory pathways (Mapk1, Hax1, Src, Fgfr3), glial responses (App, Nr3c1, C1qa) and neuronal function (Dlg4, Syngap1, Shank1, Shisa7). Remarkably, the P2 proteome showed increased levels of proteins involved in regulation of MAPK activity (Hsp90aa1, Map2k6, Dlg3), alongside proteins associated with protein dephosphorylation (Dusp14, Prkcd, Ppp6r3). Particularly, Map2k6 is a protein kinase that phosphorylates and activates p38 MAPK in response to stress; while on the contrary, Dusp14 is a protein phosphatase that inactivates p38 MAPK (80). On the other hand, the homogenate proteome also showed increased downstream protein targets of p38 MAPK such as Atf2. Notably, when active, p38 MAPK phosphorylates Atf2, which is a key transcription factor reported to play a role in regulating sleep (81). Together, our proteomic findings revealed a complex pattern of protein dysregulation associated with p38 MAPK activation. Interestingly, activation of p38 MAPK pathway has also been reported to occur in vulnerable neurons of AD patients (66).

### Impact of chronic SD on brain signaling pathways

Moreover, among its effects on post-translational modifications, SD is known to alter protein phosphorylation that might disrupt key signaling pathways in the brain and at synapses (28, 32). For instance, Guan et al. (82) reported that acute SD (6 h via exposure to novel objects) reduced phospho-ERK levels in the hippocampus, but normalized after 2 h recovery sleep, this effect was not observed in the cortex, consistent with well-established hippocampal vulnerability to sleep loss (83). Accordingly, we observed that chronic SD suppressed MAPK pathway activation in the homogenate, but not in the P2 fraction. Notably, this alteration persisted after 18 h of recovery sleep, suggesting that chronic SD induces long-lasting effects on MAPK signaling at the whole-brain level. Although previous studies have shown that acute SD (5 h via gentle handling) (60) and chronic SD (5 h/day for 28 consecutive days via gentle handling) (84) downregulates mTOR signaling in the hippocampus and the prefrontal cortex; in this study, we found no effect of chronic SD on mTOR signaling. This discrepancy suggests that region-specific SD-induced alterations in mTOR signaling could have been missed in our whole-brain analysis.

### Impact of chronic SD on vulnerable synapse ultrastructure

Chronic SD-induced changes in synaptic protein composition could have effects on synaptic structure. Numerous studies have examined the effects of acute and chronic SD on the structure of vulnerable synapses in several brain regions, using Golgi-Cox staining to measure synaptic spine density, and TEM for assessing synaptic ultrastructure (85). Indeed, Spano et al. (86) showed that acute SD (6 h via exposure to novel objects) increased synaptic density and size, but reduced the number of synaptic vesicles in hippocampal CA1 region. Interestingly, using serial block-face scanning electron microscopy and 3D reconstruction, Nagai et al. (31) demonstrated that severe SD (continuous 102 h via combination of novelty, social interactions, access to a running wheel, and other stimuli) induced ultrastructural changes in the primary motor cortex. In their study, dendritic spines containing endosomes had a larger axon–spine interface, which is an indicative of greater synaptic strength similar to wake-like state, suggesting that severe SD impairs synaptic renormalization. Moreover, Zhang et al. (87) reported that chronic SD (20 h/day for 21 days via modified multiple platform method) induced changes in the synaptic ultrastructure of hippocampal CA1 neurons, including widened and blurred synaptic spaces compared to a clearly visible synaptic cleft in controls. In our study, chronic SD altered synaptic ultrastructure of hippocampal CA1 neurons by decreasing presynapse size and synaptic vesicle density, suggesting that chronic SD impairs synaptic function and plasticity. Future electrophysiological analysis are required to determine the impact of chronic SD on synaptic transmission.

### Limitations of this study

First, evidence from humans and rodents indicates sex differences in responses to SD (88, 89). Accordingly, the use of only male mice is a limitation of this study. Future research should include female mice to determine whether these findings generalize across sexes. Second, we acknowledge that subtle chronic SD-induced transcriptional changes could have washed out in our bulk brain transcriptomic analysis due to inherent technical limitations (lack of cellular and spatial resolution). However, future transcriptomic analysis using single-nucleus RNA-seq could potentially overcome these limitations. Third, we acknowledge the limitations of using crude synaptosomes (P2 fraction) for our proteomic studies. In principle, the P2 fraction is prepared using differential centrifugation, which separates organelles based on size (51), and therefore it is inevitable to find contaminants (i.e., glial membrane fragments, free mitochondria, and myelin) (90). Importantly, the P2 fraction preparation is intended to enrich rather than purify subcellular compartments, and not intended to separate different cell types, but to separate synaptic particles from cell bodies, unbroken cells-irrespective of their identity-, large organelles such as the nucleus, and soluble proteins. Fourth, we acknowledge that 2D TEM has limitations in fully capturing the complex morphology of synapses. Future studies using advanced imaging techniques, such as 3D reconstruction and cryo-electron tomography, could provide further insights into how chronic SD alters synaptic architecture and supramolecular organization (91). Finally, this study did not assess the effects of chronic SD on cognitive performance; consequently, the transcriptomic, proteomic, signaling and synaptic ultrastructural alterations reported here cannot be directly linked to cognitive impairment.

## CONCLUSION

In conclusion, using brain transcriptomics and synaptosome proteomics, our study revealed compartment-specific protein and signaling pathway alterations induced by chronic SD, accompanied by ultrastructural alterations in vulnerable hippocampal CA1 synapses. Together our findings provide novel insights into the complex molecular dysregulation induced by chronic SD. Particularly, increased synaptic endocytosis, dendritic spine morphogenesis and long-term synaptic depression were observed in the synaptic proteome, compared with increased immune/inflammatory pathways, glial responses and neuronal function in the whole-brain proteome. Our findings support the relevance of p38 MAPK pathway, implicated in the development and progression of AD, as a mechanistic target for therapeutic interventions. Our work in WT animals represents an important foundation that can guide future work that include targeted mechanistic studies as well as studies in AD-relevant mouse models to better understand the relationships between chronic SD and AD pathophysiology.

## DATA AVAILABILITY

The data supporting the findings of this study are included within the manuscript and its associated supplementary materials (**Supplemental Table S1**). Sequencing data and proteomics data generated in this study will be available in the NCBI Gene Expression Omnibus (GEO) and the PRIDE repository.

## Supplemental Data

This article contains supplemental data.

## Acknowledgments

The authors would like to thank Dr. Ted J. Whitworth and the Center for Cellular and Molecular Imaging, Electron Microscopy Facility at Yale School of Medicine for assistance with the work presented here.

## Author contributions

C.E.-G. and S.R. conceptualization; C.E.-G., D.K., U.S., S.M., P.K., S.P., B.T., W.E.J., R.K., B.C., J.S., M.H., X.L., L.B.W., V.F., and S.R. methodology and investigation; C.E.-G., U.S., and S.R. data curation and formal analysis; S.R. funding acquisition and supervision; L.B.W. and S.R. resources. C.E.-G., U.S., S.M., and S.R. visualization; C.E.-G. and S.R. writing–original draft; C.E.-G., D.K., U.S., S.M., P.K., S.P., B.T., W.E.J., R.K., B.C., J.S., M.H., X.L., L.B.W., V.F., and S.R. writing–review and editing.

## Funding and additional information

This work was supported by the National Institutes of Health Grants R01AG071587 (S.R.), R01AG075820 (S.R. and L.B.W.), R01ES034796 (V.F.), and the Cure Alzheimer’s Fund (V.F.). This work was also supported in part by the Woodruff Faculty Fellowship at Georgia Tech (L.B.W). B.T. was funded in part by the Cell and Tissue Engineering Training Grant from the National Institute of General Medical Sciences of the National Institutes of Health under Award Number T32GM145735.

## Abbreviations

AD: Alzheimer’s disease
BP: Biological Process
CC: Cellular Component
DEGs: Differentially-expressed genes
DEPs: Differentially-enriched proteins
DIA: Data-independent acquisition
D‒PLSR: Discriminant partial least squares regression analysis
GFAP: Glial fibrillary acidic protein
GO: Gene ontology
GSVA: Gene set variation analysis
Hspa1: Heat shock 70 kDa protein 1
Iba1: Ionized calcium-binding adaptor molecule
1 LC‒MS/MS: Liquid chromatography–tandem mass spectrometry
LFQ‒MS: Label-free quantitative mass spectrometry
LV: Latent variable
MAPK: Mitogen-activated protein kinases
MF: Molecular Function
mTOR: Mammalian target of rapamycin
NfL: Neurofilament light chain
PCA: Principal-component analysis
PSD: Postsynaptic density
PSD95: Postsynaptic density protein 95
RT: Room temperature
SD: Sleep disruption
SDHA: Succinate dehydrogenase complex flavoprotein subunit A\
SEM: Standard error of the mean
STAT1: Signal transducer and activator of transcription 1
SVs: Synaptic vesicles
SV2A: Synaptic vesicle glycoprotein 2A
TEAB: Triethylammonium bicarbonate
TEM: Transmission electron microscopy
TOMM20: Translocase of outer mitochondrial membrane 20
UPR: Unfolded protein response
VDAC: Voltage-dependent anion channel
WT: Wild-type
ZT: Zeitgeber time

